# A multivariate view of cognitive performance reveals positive correlation in the Trinidadian Guppy (*Poecilia reticulata*)

**DOI:** 10.1101/2021.11.04.467320

**Authors:** Pamela M Prentice, Alex Thorton, Alastair J Wilson

**Affiliations:** Centre for Ecology and Conservation, University of Exeter, Penryn Campus, Cornwall, TR10 9FE, UK

**Keywords:** Cognitive performance, among-individual variation, cognitive evolution, multivariate mixed model, correlation matrix, Trinidadian guppy

## Abstract

Cognitive variation is common among-individuals and can be consistent across time and context. From an evolutionary perspective, among-individual variation is important as a pre-requisite for natural selection and adaptive evolution. Selection is widely hypothesized to favor high cognitive performance but directional selection should erode variation over time, how then is cognitive variation maintained? As selection does not act on traits in isolation, covariance among specific cognitive traits and/or other aspects of phenotype (e.g. personality) could result in fitness trade-offs that are important in shaping evolutionary dynamics. Here we test this using Trinidadian guppies (*Poecilia reticulata*), using a multivariate approach by characterising the correlation structure among task-specific cognitive performance measures and a personality trait. We estimate the among-individual correlation matrix (**ID**) in performance across three cognitive tasks; colour association learning task; motor learning task; reversal learning task, and the personality trait ‘boldness’ measured as emergence time from a shelter. We found no support for trade-offs among performance in these tasks. Nor do we find evidence of hypothesised speed-accuracy trade-offs within the association learning task. Rather we find strong positive correlation structure in **ID,** with 57% of variation explained by the leading eigen vector. While noting that non-cognitive factors and assay composition may affect the structure of **ID**, we suggest our findings are consistent with the *g*-model of cognitive performance variation, in which a dominant axis of variation loads positively on all performance measures. Thus, we add to a growing body of support for general variation among individuals in animal cognitive ability.

## Introduction

Interest in the cognitive mechanisms by which animals acquire, process, store and use information from the environment (Healy & Rowe, 2010; Shettleworth, 2010), has grown substantially in recent years among ecologists and evolutionary biologists. Differences in cognitive abilities between species have long been recognised (Wasserman & Zentall, 2006; Shettleworth, 2009), but we now know that individuals within a species can differ in cognitive performance in a manner that is consistent across time and context (for a review, see Cauchoix et al., 2018, but see Troisi et al., 2021 for an exception to this). From an evolutionary perspective, variation at this level is required for natural selection and is thus central to our understanding of adaptive evolutionary dynamics (Roff, 2002). However, how or why this variation is maintained within populations is not always clear. For instance, while we might intuitively expect selection to favour high cognitive performance, directional selection of this form is generally expected to erode variation over time. One possibility is that fitness trade-offs could shape patterns of variation in cognition, just as they do in other aspects of phenotype (e.g. life history; Stearns, 1992). If so, do trade-offs arise among different cognitive ‘domains’ or traits, with variation maintained because individuals (or genotypes) that perform better than average in some respects, perform worse in others? Here we test this idea using a multivariate approach to characterise and test hypothesised relationships of cognitive performance between multiple variables measured within three cognitive tasks using the Trinidadian guppy (*Poecilia reticulata*). We ask whether there is support for negative correlation structure, as expected under the trade-off hypothesis, or whether in fact correlations among performance measures are uniformly positive – a pattern predicted if multivariate cognitive variation is explained by a single general cognitive performance factor or “general intelligence” (*g*) (Nisbett et al., 2012).

Many empirical studies have now shown that animal populations can harbour high levels of variation in cognitive performance among individuals (Boogert et al., 2018), and this has been evidenced across different taxonomic groups, including insects (e.g. Li et al., 2017), fish (e.g. Lucon-Xiccato & Bisazza, 2017), birds (e.g. Ashton et al., 2018) and mammals (e.g. Mazza et al., 2019). Among-individual variation is also found across different aspect of cognition (e.g. spatial memory (Sonnenberg et al., 2019); association learning (Kniel et al., 2020), inhibitory control (Brandão et al., 2019)). Although numerous studies have explored relationships between cognitive performance in different cognitive tasks (for a review see Burkart et al., 2017; Boogert et al., 2018), relatively few studies have characterised the among-individual covariance (or correlation) structure between these. This is important to do because trade-offs between domains have been hypothesised to maintain cognitive variation at the individual level (Del Giudice & Crespi, 2018). This hypothesis leads to an expectation of negative correlation structure which is exemplified by predictions of ‘trade-offs’ in performance across different cognitive domains (Shettleworth, 2012; Thornton & Lukas, 2012).

It has been hypothesised that trade-offs, such as those observed between learning and cognitive ‘flexibility’ (Del Giudice & Crespi, 2018), could maintain variation in performance. Flexibility is broadly defined as the ability to adapt when environmental stimuli or information cues change. It is often tested by reversal learning experimental paradigms in which individuals must be flexible to override old cue-reward associations and form new ones (Bitterman, 1965; Kehagia et al., 2010; Buechel et al., 2018) . For example, in Florida scrub-jays (*Aphelocoma coerulescens*) individuals that were quick to learn an initial colour-reward cue (high associative learning performance), were slower to adjust when the cue signal was reversed (Bebus et al., 2016). The implication of such patterns is that some individuals are more ‘intrinsically driven’ than others; they learn initial associations quickly but, being less sensitive to external stimuli, struggle to adapt when cues are altered. However, counter-examples can be found. For instance, those individual bumblebees (*Bombus terrestris*) that quickly learned to discriminate between two colours (where one was associated with a floral reward), were also faster to learn a new association when the cues were reversed (Raine & Chittka, 2012).

Trade-offs have also been postulated to maintain variation in performance within cognitive domains. Speed-accuracy trade-offs are perhaps the best known example (Chittka et al., 2009). Faster decision-making should provide a competitive advantage (e.g. by increasing the potential rate of resource acquisition), but by allowing less time to assess environmental cues, error rates may be increased. In contrast, slower decisions may be more accurate, but being slow to act can mean resources are lost to competitors. A number of empirical studies have provided support for speed-accuracy trade-offs. For example, individual archerfish (*Toxotes chatareus*) that tended to make slower decisions within a colour discrimination task also had greater accuracy (Jones et al., 2020). A similar result was found in zebrafish (*Danio rerio*) given a visual discrimination task (Wang et al., 2015), and in wild-caught great tits (*Parus major*) given a foraging task (Moiron et al., 2016). However, once again counter examples also exist. For example, a prior study of guppies found no support for a speed-accuracy trade-off using a shape and a colour discrimination task (Lucon-Xiccato et al., 2016), and three-spined sticklebacks (*Gasterosteus aculeatus*) making fast decisions in a spatial leaning task did not show reduced accuracy (Mamuneas et al., 2015).

Thus, while trade-offs between and within components of ‘multivariate’ cognitive ability are intuitive and evidenced in some cases, they may not be inevitable and positive correlations between traits and domains are also found (Guenther & Brust, 2017; Wallace & Hofmann, 2021). If correlation structure among cognitive performance traits is universally positive (with respect to expected fitness consequences), then variation among individuals can be explained by a domain-general structure that invokes a single latent general cognitive performance or “intelligence” factor, sometimes denoted ‘*g*’ (Deaner et al. 2006; Lefebvre and Sol 2008). While there is evidence for this in humans (Deary et al., 2010; Burkart et al., 2017), support for the *g*-model in non-human animals systems remains limited (but see (but see Hopkins et al., 2014; Shaw et al., 2015; Arden & Adams, 2016; Ashton et al., 2018). Note that, under this model, trade-offs may still be important for maintaining cognitive variation, but if so they must operate between general cognitive performance and other (non-cognitive) aspects of phenotype. This possibility is mirrored elsewhere in the behavioural literature since for example, individual personality can be viewed as a component of an extended ‘life history’ (Sih et al., 2004; Wolf et al., 2007). Indeed, given that personality traits and cognitive performance are frequently correlated (Guenther et al., 2014; Nawroth et al., 2017; White et al., 2017), and both thought to drive variation in resource acquisition, then trade-offs between them are plausible (Sih & Del Giudice, 2012; Dougherty & Guillette, 2018).

Here, we investigate among-individual (co)variation between multiple measures of cognitive performance variables across three cognitive tasks (associative learning task, motor learning task and a reversal learning task) in a captive population of wild-type guppies (*Poecilia reticulata*). Guppies have been widely used as a model in behavioural and evolutionary ecology, and methods for assaying among-individual variation in cognitive performance are well established (Laland & Reader, 1999; Miletto Petrazzini & Agrillo, 2016). This species is known to perform well (on average) in tests of associative learning (Lucon-Xiccato & Bisazza, 2016; Kniel et al., 2020), motor learning (Lucon-Xiccato & Bisazza, 2016; Lucon-Xiccato et al., 2017), and cognitive flexibility (Cauchoix et al., 2018; Fong et al., 2019). However, we know little about if and how cognitive performance measures within these distinct tasks covary among individuals. Broadly stated, our goal is to estimate the structure of this multivariate variation to evaluate whether trade-offs among cognitive performance measures are evident, or whether among-fish variation is consistent with general cognitive performance (*g*). We also seek to ask whether trade-offs shape variation in cognitive performance within cognitive tasks, and whether personality variation is linked to, or trades off against cognitive performance variation in this species.

We note that the extent to which among-trait associations shape, and in the particular context of trade-offs, constrain evolutionary adaptation strictly depends not on the phenotypic correlation structure, but on the genetic contribution to this. In particular, evolutionary constraint arises from the genetic (co)variance structure (**G**) and its alignment (of lack thereof) with selection (Jones et al., 2004; Lande, 1979; Walsh et al., 2009). Unfortunately estimation of **G** is challenging in general (requiring large volumes of data from related individuals; Wilson et al., 2010), and particularly so for cognitive performance measures. This is because high-throughput phenotyping of cognitive performance across multiple tasks is notoriously laborious and challenging. Consequently, very few studies have estimated **G** for sets of cognitive performance measures in non-human animals (but see Langley et al., 2020). Thus most multivariate studies of cognition, and behavioural phenotypes more generally, rely on Cheverud’s conjecture (Cheverud, 1988) that phenotypic patterns of covariation can be used to infer evolutionarily important relationships between traits. Accepting this view uncritically implies that the (co)variance structure of the phenotypic matrix **P** (historically referred to as the phenotypic gambit; Grafen, 1984), which can be readily estimated in a set of individuals observed once only for each of the target traits, can yield robust insights into the structure of **G**. However, behavioural traits and cognitive performance measures are typically very plastic and often subject to high levels of measurement error (e.g. relative to morphological traits), in addition to sources of environmental variation. These phenomena lead to high levels of within-individual variation such that a single observation may tell us relatively little about an individual’s phenotype (and so genotype) (Brommer, 2013). In other words, the extent to which **P** is a good proxy for **G** will decline if **P** is dominated by within-individual variation (plasticity, measurement error) versus among-individual variation (which includes genetic factors).

In the present study we therefore adopt an intermediate approach in which we estimate the among-individual (co)variance matrix (**ID**) rather than either **G** or **P**. We do this by obtaining repeated measures of performance on individual fish. Since **G** is a component of **ID,** while **ID** is a component of **P**, all else being equal we expect the among-individual covariance matrix to be better proxy of the genetic matrix than the total phenotypic matrix. We target multiple cognitive performance measures across three cognitive tasks (association learning task, motor learning task and a reversal learning task) allowing us to: 1) test whether individuals differ in their cognitive performance between distinct cognitive tasks. We predict that individuals will differ in all measures of cognitive performance within all three tasks; the association task, the novel motor learning task, and the reversal learning task as well as in their rates of improvement with experience (interpretable as learning). We then estimate the **ID**- matrix among all cognitive performance measures across the three different cognitive tasks, in addition to a shy-bold type personality trait (henceforth referred to as ‘boldness’). We ask 2) whether the **ID**- matrix provides evidence for trade-offs among cognitive performance measures (exemplified as a negative correlation structure in the **ID**-matrix) or whether it is dominated by positive correlations consistent with the *g*-model of general cognitive performance; 3) whether speed accuracy trade-offs are present within both the association task and reversal learning task; and finally 4) whether among-individual variation in shy-bold personality trait ‘boldness’ is correlated with cognitive performance variation in the three cognitive tasks.

## Methods

### Ethics

This work was conducted under the auspices of the Animals (Scientific Procedures Act) with approval of the University of Exeter research ethics committee, under licence from the Home Office (UK) (Licence Number PPL30/3256). Experimental procedures and behavioural assays were developed in accordance with the principles of the three Rs and ASAB guidelines (Buchanan et al., 2020). All periods of handling and emersion were kept to a minimum and only fish deemed healthy and exhibiting normal behaviour were used in trials. At the end of the experiment, fish were returned to a designated ‘retirement’ tank (containing females as well as males) and not used in any further experiments.

### Husbandry

All behavioural assays were carried out between October 2019 and January 2020 inclusive. Data was collected from captive-bred guppies bred and housed at the fish laboratory at the University of Exeter’s Penryn campus. The population is descended from wild fish caught in February 2017 from the lower Aripo River, Trinidad and has been subsequently maintained with no deliberate selection or inbreeding. All fish housed in the laboratory were fed to satiation twice daily (0800 – 1000h and again at 1600 – 1800h) using commercial flake food and live *Artemia nauplii*. Water temperature was maintained at 23-24°C in well-aerated closed system tank stacks that undergo 25% water changes each week and with weekly tests for ammonia, nitrate and nitrite levels. Lighting was kept at a 12:12 light/dark cycle.

### Experimental apparatus

Adult males (n = 43) and females (n = 37) were sampled from the stock population. Sampling was random but we approximately size matched fish within each sex (e.g. by avoiding very large females). Fish were then housed singly in separate tanks (15 l, 18.5 × 37 × 22cm) for the duration of behavioural testing. The tank set up closely followed that used by Lucon-Xiccato & Bisazza (2014), each being divided equally into a ‘home’ compartment at the rear of the tank, and a ‘test’ compartment at the front of the tank, separated into two compartments (using white plastic) by a guillotine door (Figure 1a). The rear ‘home’ compartment (20 x 18.5 cm) allowed individuals visual access to fish in neighbouring tanks and contained foliage to provide coverage and environmental enrichment (Brydges & Braithwaite, 2009). Conversely, the ‘test’ compartment (17 x 18.5 cm) was screened from neighbours (using white plastic) to prevent any possibility of social learning (i.e. by observation of neighbours) influencing cognitive task performance. The ‘test’ compartment contained a white plastic plate (4 x 10 cm) placed on the gravel substrate, perforated with 2 equally spaced wells (Figure 1b). A total of 48 experimental tanks were used. These were contained within two ‘stacks’, each comprising 24 tanks (8 tanks per row, 3 rows high) on a shared recirculating water supply. As the testing protocol (described below) took 18 days per fish, data were in practice collected in 3 ‘blocks’ over a total testing period spanning 12 weeks. Before cognitive testing all fish were allowed to acclimate to experimental tanks for 48 hours. During this period they were fed twice daily with bloodworm (*Chironomidae* larvae) pipetted into one (randomly chosen) well of the white plastic plate and the guillotine door was left open allowing unrestricted use of both compartments.

**Figure 1.**
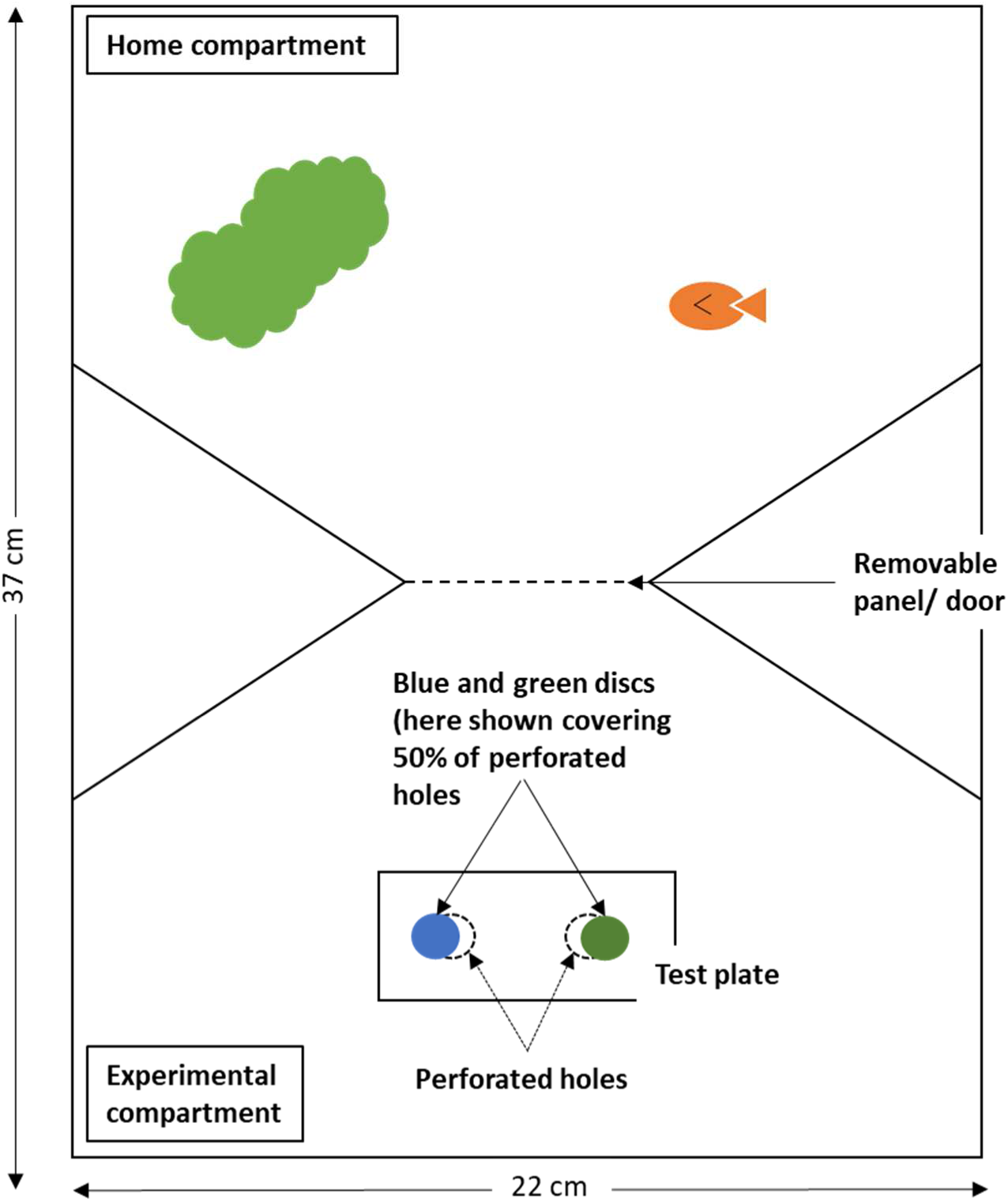
Aerial view of the tank used to house individual fish throughout the experiment. The home compartment allowed the focal fish visual access to neighbouring tanks and included foliage plants for cover and enrichment. Also shown in the experimental compartment is the test plate used during each task.

### Cognitive assay

We used a repeated measures design to test multiple cognitive performance measures across three different cognitive tasks within a single, extended, testing paradigm. Each fish was observed up to 63 times as it emerged from the home compartment to obtain a food item that had been placed in one of two wells in the test compartment. Individual fish were trained to discriminate between a rewarded and an unrewarded colour cue, indicating which well the reward could be obtained from. In all trials, food items (bloodworm) were placed in both wells, but access was restricted by a small plastic disc in the well with the unrewarded colour cue. This was to ensure that olfactory cues would be insufficient to locate available food, instead fish were required to learn to associate a specific colour with reward access. The rewarded colour (either blue or green) was randomly determined for each fish prior to testing, ensuring balance across subjects to control for any pre-existing colour bias. The relative position of the rewarded well (left versus right) for each trial was randomised and we also controlled for any effect this could have in our statistical modelling (described later).

The 63 observations per individual were grouped into 7 ‘sets’ of 9 ‘trials’ (observations). Each set spanned a 2 day period, which was followed by a rest day before the next set. Test conditions were altered between trial sets, by adjusting the position of green and blue coloured plastic counters. Specifically, after each set, the association task increased in difficulty as the coloured counter covered the holes to an increasing extent. This also allowed extraction of information on different cognitive traits as we now describe. In set 1, the coloured counters covered 0% of the wells, in set 2 we increased this to 25%, then subsequently to 50% (set 3), 75% (set 4) and 100% (set 5). Once the counters covered 75% of the wells (set 4), guppies had to dislodge the counters in order to reach the reward in the well underneath. We consider this a novel motor learning task, as fish had to learn to dislodge the counter by physical manipulation. The difficulty of this task is increased with 100% coverage (set 5). In the final two sets (set 6 and 7) of 9 trials we conducted a reversal learning task, by reversing the colour cue -food reward association previously learnt in the first 5 sets.

Prior to each trial, fish were guided into the ‘home’ compartment of the tank with a net, and the guillotine door was closed. The experimenter set up the plate in the ‘test’ compartment, by pipetting bloodworm into the wells and the two coloured counters placed either to the side of (set 1), or covering the rewarded well (partially or fully depending on set number as described). The food in the unrewarded hole was covered completely by a small plastic disc. The trial started once the door to the experimental compartment was opened and continued until the fish ate the bloodworm from the rewarded well, or until 10 minutes, whichever occurred first (see the electronic supplementary material, video S1). All trials were recorded with a GoPro Hero 6 camera, mounted in front of the tank and behavioural scoring of videos was quantified using the software BORIS (Friard & Gamba, 2016). We imposed a deliberately weak learning criterion, removing any fish that failed to achieve a minimum of 5/9 successes (defined as locating and/or eating the food reward) at the end of each set. In fact, we would have ideally avoided selective removal of fish such that among-individual variation seen in later sets can be interpreted as fully representative of the initial population sampled. However, it is also the case that estimated correlations between task-specific performance may be misleading if the ability to solve each task is closely associated with the number of prior learning opportunities (Guez & Griffin, 2016; Shaw & Schmelz, 2017), which in turn depends on differences in, for example, motivation or neophobia. Furthermore, since additional food (beyond the reward items) was not provided during the behavioural testing, the removal strategy adopted ensured that any individuals not engaging with the task at all could not be food deprived for more than 48 hours. Over the course of the data collection, 16 individuals were removed before the end of set 6.

### Cognitive performance measure definitions

Data obtained from the trials were used to define observed proxies of traits, including one personality trait (*boldness*) and six measures of cognitive performance that were collected during three distinct cognitive tasks (*associative learning, motor learning,* and *reversal learning*). In some cases cognitive performance measures were defined during trials in a single set (9 trials in total), but where data from 2 sets are equally relevant (18 trials in total); these were combined to maximise informative observation number and hence maximise statistical power. The specific personality trait and cognitive performance measures are described below and are summarised for reference in Table 1.

**Table 1.** Outline of behaviours used to assess performance within each measure of personality and cognitive performance, including details of corresponding set number at which each variable was measured and corresponding acronyms used within the text.

| Set Number/<br>Number of<br>informative trials | Personality trait & Cognitive<br>performance task / Variable<br>measured | Description of variable measured | Acronym |
| --- | --- | --- | --- |
| Set 1, 9 trials | Boldness | Time to emerge<br>(Time taken to emerge into the test compartment once the compartment door is opened) | *emergence* |
| **Association learning task** |  |  |  |
| Set 1, 9 trials | Decision speed | (After emergence into test compartment, time taken to make first choice, measured as swimming within 1 body length of, and actively exploring correct counter.) | *AL<sub>speed</sub>* |
| Set 1, 9 trials | Decision accuracy | (Accuracy of first choice. Either correct or incorrect) | *AL<sub>accuracy</sub>* |
| Set 2 & 3, 18 trials | Time to eat | (After emergence into test compartment, time taken to eat reward from the correct well) | *AL<sub>time</sub>* |
| **Motor learning task** |  |  |  |
| Set 4 & 5, 18 trials | Time to eat | (After correct choice made (i.e. swimming within 1 body length of, and actively exploring correct counter), time taken to dislodge counter and eat the food reward) | *ML<sub>time</sub>* |
| **Reversal learning task** |  |  |  |
| Set 6 & 7, 18 trials | Time to eat | (After emergence into test compartment, time taken to dislodge disc and eat reward from the correct well) | *RL<sub>time</sub>* |
| Set 6 & 7, 18 trials | Decision accuracy | (Accuracy of first choice. Either correct or incorrect) | *RL<sub>accuracy</sub>* |

#### Boldness

During the first 9 trials (set 1), time taken to emerge from the home compartment into the experimental compartment was recorded (*emergence*), as a proxy of consistent differences in personality trait ‘boldness’ (White et al., 2016, 2020).

#### Associative learning

We focused on the first 9 trials (set 1) to characterise associative learning and test for speed-accuracy trade-offs. A similar study to the current testing paradigm, guppies learnt a colour association in 9.18+2.50 trials (Bisazza et al., 2014), as such we expected that (on average) fish would have effectively learnt the association by the end of set 1 such that any speed accuracy trade-off would be less relevant to observed outcomes in later sets. We recorded three cognitive performance measures in each of the 9 trials in set 1. The first measure was whether the first counter explored in each trial was the correct (1) or incorrect (0) choice for obtaining the reward (*AL_accuracy_)*. A choice was deemed to have been made if individual fish swam within 1 body length of, and was actively exploring a counter (with head and body directed towards the stimulus counter). We note that accuracy of the first counter explored in trial 1 of set 1 will be by chance alone due to the absence of prior experience. The second cognitive performance measure was time taken to make this first choice in each trial (*AL_speed_*), measured in seconds from emergence into the test compartment. To evaluate overall performance in association learning, we used total time taken from emergence to find and eat the food reward in each trial (*AL_time_*). Note, when modelling this variable we maximised observation number (and hence statistical power) by using observations from 18 informative trials; 9 trials in set 2 and 9 trials in set 3. During set 2 and 3, the counters covered 25% and 50% of the wells respectively. This level of coverage still allowed fish to access the reward without any physical manipulation of the counter.

#### Motor Learning

To assess performance in the novel motor learning task, we used observations from 18 informative trials; 9 trials in set 4 and 9 trials in 5 (where the counter covered 75% and 100% of the well area respectively). Note that at 75% coverage fish were required to physically manipulate the counter in order to reach the food reward. We used time taken by each individual in all trials to obtain and eat the food reward (*ML_time_*), measured as time taken to obtain the food reward after they had made the correct choice of counter in each trial.

#### Reversal learning

This was assessed using observations from 18 informative trials: 9 trials in set 6, and 9 trials in set 7. During these trials the reward colour was reversed. Note that counter coverage of the wells remained at 100% and therefore fish required to dislodge the counter in order to access the reward. For this reversal learning task we recorded the accuracy of each individual’s first choice in every trial in set 6 and 7, (*RL_accuracy_)* as well as the total time taken, from emergence into the test compartment, to locate and eat the food reward in each trial (*RL_time_*).

### Statistical analysis

We used univariate and multivariate linear mixed effect models to characterise among-individual variation in and covariation among the personality trait and cognitive performance measures defined. For analysis, the observed (censored) time variables from the cognitive tasks were natural log-transformed (*AL_speed_, AL_time_, ML_time_* and *RL_time_*), and then all continuous time variables were multiplied by -1 (*boldness*, *AL_speed_, AL_time_, ML_time_* and *RL_time_*). Following this, all response variables were mean centred and scaled to standard deviation units. The log transformation was to improve the assumption of Gaussian error structure, multiplication by -1 was just to ease of interpretation (making larger values equate to higher performance). Scaling to standard deviation units was also to simplify interpretation of estimated variance components (explained further below). All models were fit by restricted maximum likelihood (REML) using ASReml-R 4.1 (Gilmour et al., 2009; Butler et al., 2018), R version 3.6.1 (R Core Team, 2017). We make the standard assumptions that random effects and residuals are normally distributed with means of zero and variances to be estimated. We acknowledge that these assumptions are necessarily violated since (i) the two accuracy traits are recorded as binary variables and (ii) latency (time) traits are censored at 10 minutes. However, we consider this strategy both pragmatic and justifiable given that Gaussian mixed models are largely robust to violations of distributional assumptions (Schielzeth et al., 2020) while (Bayesian) multivariate generalised mixed models remain challenging to implement.

#### Univariate analysis

First, we fit separate univariate mixed models to test for and characterise among-individual variation in each of our 7 traits (*boldness*, *AL_speed_, AL_accuracy_, AL_time_, ML_time_, RL_time_* and *RL_accuracy_)*. All models included a fixed effect of the mean, as well as fixed factors of *sex*, *stack* (denoting which of two aquaria stacks the fish was tested in), *colour* (denoting the colour of counter individual fish were trained to associate with the rewarded cue; green or blue), and *reward side* (denoting which side the rewarded counter was placed on; left or right). We also included *trial number* (i.e. the repeat number of trial, fitted as a factor to avoid assuming a linear functional form for any change in the mean across repeats). For each trait, we firstly fit this ‘null’ model without random effects, and then refitted with *individual identity* as a random intercept. For each trait, we compared the full (i.e. with *individual identity* included) and null models by likelihood ratio tests (LRT) to obtain a statistical test of the among-individual variance (V_I_). To test a single variance component (which cannot be less than zero) we assume twice the difference in log-likelihood between the full and reduced models is distributed as a 50:50 mix of χ^2^_0,1_ and χ^2^ as recommended by Visscher (2006). Since transformed traits are analysed in standard deviation units, total observed variance is 1 and V_I_ can actually be interpreted as an estimate of repeatability. However, we also calculated estimates of the adjusted repeatability (R) which is the proportion of phenotypic variance conditional on fixed effects that is explained by among-individual differences (Nakagawa & Schielzeth, 2010). Thus R = V_I_ / V_P_ where V_I_ is the among-individual variance and V_P_ is sum of V_I_ and V_R_, the residual or within-individual variance. Conditional F-statistics were used to determine the significance of fixed effects in each (full) model, and we elected not to perform model simplification, as we wanted repeatability estimates to be conditioned on a common set of fixed effects and thus comparable across abilities.

#### Multivariate analysis

To test for the presence of, and investigate the structure of, the (co)variance **ID**-matrix, we built a multivariate mixed model among all cognitive performance traits and ‘boldness’. All response variables were transformed and scaled as per the univariate modelling procedure above, and we note that multiplication by -1 of all the observed (censored) time traits simplified interpretation of the correlation structure (as larger values equate to higher performance in continuous traits). Fixed and random effects were fitted on each trait as specified for the univariate models. Fixed effects estimates are reported in the supplementary information (Table S1). We specified **ID** as an unstructured matrix to be estimated. Note that observation-level (residual) covariances are not statistically identifiable if traits are not measured at the same time. We thus specified a residual covariance matrix structure in which covariances between traits measured in different sets were fixed to zero. We tested for overall among-individual covariance among the traits by comparing this model against a reduced one in which **ID** was specified as a diagonal matrix (i.e., among-individual variances are estimated but covariances are assumed to equal zero).

To aid biological interpretation of the **ID**-matrix, we rescaled our estimate to the corresponding correlation matrix (noting that for any pair of traits (x,y), r_I(x,y)_ = COV_I(x,y)_/ (V_I(x)_V_I(y)_)^0.5^). We further scrutinised the correlation (r) structure among cognitive (c) traits only using **ID_rc_**, defined as the 6x6 submatrix of **ID** obtained after putting on a correlation scale and dropping the row/column corresponding to boldness. **ID_rc_** was subject to eigen vector decomposition to provide a descriptive view of the major axes (or principle components) of variation. Under the *g* model, we would expect uniformly positive correlation structure in **ID_rc_** which would be reflected by a dominant first eigen vector loaded on all traits with the same sign. Conversely, if trade-offs dominate the structure of among-individual variation in the multivariate cognitive phenotype **ID_rc_** should contain at least some negative correlations and one or more important principle components with loadings that are antagonistic in sign between traits in trade-offs. We estimated approximate 95% confidence intervals on the eigen values (scaled to proportions of variation in the matrix) and on the trait loadings associated with each principal component using a parametric bootstrap approach described in Boulton et al (2014).

## Results

Visualisation of the raw data shows a varied effect of trial number on the personality trait ‘boldness’, and all performance measures across the cognitive tasks (Figure 2; see Supplemental Information Figure S1 a–g of raw data across all sets of trials). Patterns of improvement are observed in the raw data for the following: average time to *emergence* (boldness; Fig 2a); *AL_speed_* (Fig 2b) and *AL_accuracy_* (Fig 2c) measured within the first set of trials of the association learning task; and both *RL_time_* (Fig 2f) and *RL_accuracy_* (Fig 2g) measured within the reversal learning task (although the trend in the former is weak). Time-based measures of cognitive performance for *AL_time_* and *ML_time_* did not uniformly improve (i.e. reduce) over the sets of trials used to assay them, as might perhaps be intuitively expected (Figure 2). Note however, that (i) we deliberately chose to use *AL_time_* recorded from sets 2 and 3 to capture variation in the extent to which the association had been learned after the experience of set 1 (during which rapid improvement in mean performance is seen as predicted; Fig S1d). Additionally, (ii) both *AL_time_* and *ML_time_* were assayed using 2 consecutive sets of 9 trials separated by an increase in task difficulty (i.e., change in position of the counter between sets) imposed between trial 9 and 10. In fact *ML_time_* does show a general pattern of improvement within each set separately (sets 4 and 5; Supplemental Information Fig S1e) that is not apparent from the linear trend across all 18 trials (Fig 1e).

**Figure 2.**
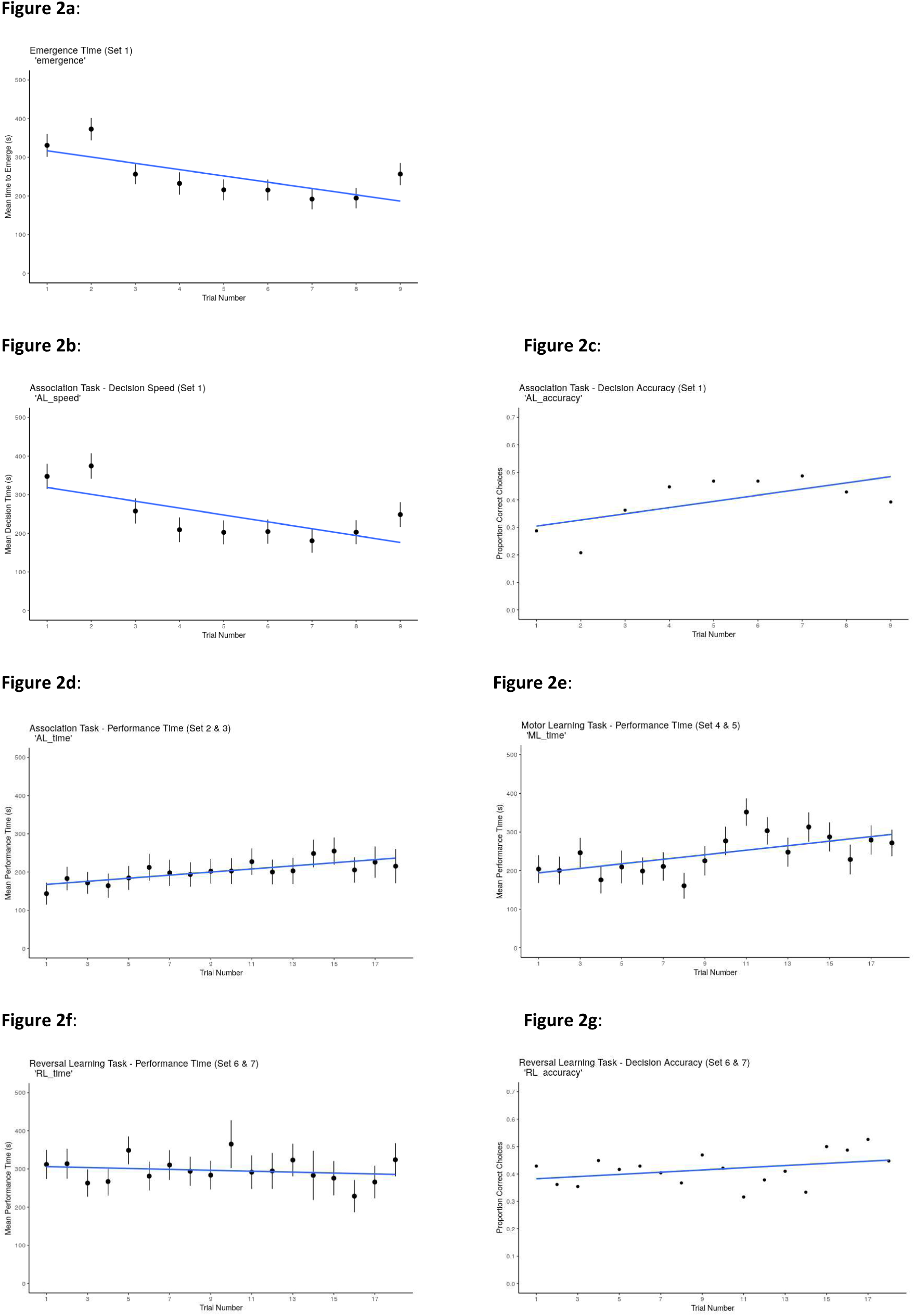
Plots of raw data of emergence data and cognitive performance measures. Each plot header outlines the set numbers used to assay each variable; a) emergence time (*emergence)*; association learning task b) decision speed (*AL_speed_*) and c) accuracy (*AL_accuracy_*), d) association learning task performance time *(AL_time_*); e) *_motor_* learning task performance time (*ML_time_*); f) reversal learning task performance time (*RL_time_*) and g) accuracy (*RL_accuracy_*). For continuous variables (*AL_time,_ AL_speed_*, *ML_time_* and *RL_time_*), higher values (seconds) represent longer durations to locate the food reward in each task, and lower values represent faster durations to solve the task. For binary variables (*AL_accuracy_* and *RL_accuracy_*), accuracy is represented as the percentage of correct choices made in each trial. Error bars represent mean and standard errors of performance time for individuals. Blue line represents regression line through mean performance or accuracy at each trial.

The univariate mixed models confirmed the statistical significance of these qualitative patterns in the raw data with respect to changes in mean performance across trials (see Supplementary Information Table S1 *a–g)*. Noting that models were run on transformed data such that positive effects always equate to an increase in performance. Furthermore, *trial number* was fitted as a factor to avoid assuming linear trends. Noting these points, we found significant effects on the following: *emergence* (F_8, 555.7_=9.24, P< 0.001; with coefficients becoming more positive as *trial number* increased; Table S1)*, AL_speed_* (F_8, 555.0_=7.380, P< 0.001; with coefficients becoming more positive)*, AL_accuracy_* (F_8, 555.0_=4.602, P< 0.001; with coefficients becoming more positive), *AL_time_* (F_17, 801.9_=2.246, P=0.002; with coefficients becoming more negative, indicating a decrease in performance), and *ML_time_* (F_17,692.6_=1.174, P< 0.001; coefficients becoming more negative, indicating a decrease in performance). The effect of *trial number* on (mean) performance was not significant in *RL_time_* (F_17, 581.2_=1.565, P= 0.060) and *RL_accuracy_* (F_17, 584.2_=1.085, P= 0.271). The significance and magnitude of other fixed effects included is not directly relevant to any hypotheses being tested and we do not discuss them further, but they are reported in full in the supplemental materials (see Supplementary Information Table S1).

With respect to among-individual variation, likelihood ratio tests (LRT) on univariate mixed models provided strong statistical support for the presence of among-individual variance in all seven traits tested (LRT of model 1 vs. model 0; all *P* < 0.001; Table2). Repeatabilies (R) estimated conditional on fixed effects (i.e. as R = V_I_/V_P_ = V_I_/(V_I_+V_R_)) were low to moderate, ranging from 20-46% with a median across the seven traits of 33.5% (Table 2). As well as among-individual variance, the multivariate modelling shows strong support for among-individual covariance in **ID**. Likelihood ratio comparison of the full model to a reduced fit in which all cross-trait covariances in **ID** were set to zero showed the former was a significantly better fit to the data (χ2_21_=242.756, P<0.001). Among variables measured, the estimated covariances were positive between all trait pairs (Table 3). Boldness, as captured by *emergence* was strongly (and significantly; based on assuming approximate 95% CI of r_I_ ±1.96 SE) correlated with cognitive performance as measured by all variables in the association learning task ((r ± (SE)), *AL_time_* = 0.663 (0.095), *AL_speed_* = 0.818 (0.046), *AL_accuracy_* = 0.788 (0.062); Table 2), and to decision accuracy within the reversal learning task (*RL_accuracy_* = 0.412 (0.171)).

**Table 2:** Estimated among-individual variance (V_I_) and adjusted repeatability (R) from univariate models of all cognitive performance and personality measure. Standard errors are shown in parentheses. Also shown are likelihood ratio tests of V_I_.

| Task | Variable | $V_i$ | $R$ | $\chi^2_{0,1}$ | $P$ |
| --- | --- | --- | --- | --- | --- |
| *Associative Learning* | *Emergence* | 0.415 (0.077) | 0.439 (0.049) | 206.004 | <0.001 |
|  | *AL<sub>speed</sub>* | 0.442 (0.081) | 0.460 (0.049) | 220.393 | <0.001 |
|  | *AL<sub>accuracy</sub>* | 0.202 (0.045) | 0.264 (0.046) | 79.989 | <0.001 |
|  | *AL<sub>time</sub>* | 0.341 (0.074) | 0.335 (0.050) | 186.345 | <0.001 |
| *Motor Learning* | *ML<sub>time</sub>* | 0.275 (0.065) | 0.329 (0.054) | 171.462 | <0.001 |
| *Reversal Learning* | *RL<sub>time</sub>* | 0.372 (0.088) | 0.375 (0.058) | 178.426 | <0.001 |
|  | *RL<sub>accuracy</sub>* | 0.182 (0.050) | 0.203 (0.047) | 65.327 | <0.001 |

**Table 3.** ID. variance–covariance–correlation matrix from the multivariate mixed model. Estimated variances are shown on the diagonal (dark grey shading), with correlations above and covariances below. Standard errors are shown in parentheses, and bold font denotes nominally significant estimates assuming approximate 95% CI of ± 1.96SE.

| ID Matrix | *emergence* | *AL<sub>speed</sub>* | *AL<sub>accuracy</sub>* | *AL<sub>time</sub>* | *ML<sub>time</sub>* | *RL<sub>time</sub>* | *RL<sub>accuracy</sub>* |
| --- | --- | --- | --- | --- | --- | --- | --- |
| *emergence* | **0.426 (0.0773)** | **0.818 (0.046)** | **0.788 (0.062)** | **0.663 (0.095)** | 0.239 (0.164) | 0.166 (0.176) | **0.412 (0.171)** |
| *AL<sub>speed</sub>* | **0.361 (0.072)** | **0.458 (0.082)** | **0.894 (0.039)** | **0.758 (0.074)** | 0.271 (0.155) | 0.228 (0.169) | **0.397 (0.168)** |
| *AL<sub>accuracy</sub>* | **0.242 (0.052)** | **0.285 (0.056)** | **0.221 (0.046)** | **0.729 (0.092)** | 0.124 (0.175) | 0.090 (0.184) | 0.258 (0.190) |
| *AL<sub>time</sub>* | **0.286 (0.072)** | **0.340 (0.076)** | **0.227 (0.055)** | **0.437 (0.092)** | **0.747 (0.082)** | **0.540 (0.129)** | **0.423 (0.161)** |
| *ML<sub>time</sub>* | 0.092 (0.067) | 0.108 (0.067) | 0.034 (0.049) | **0.291 (0.075)** | **0.346 (0.078)** | **0.817 (0.067)** | **0.340 (0.167)** |
| *RL<sub>time</sub>* | 0.074 (0.082) | 0.106 (0.083) | 0.029 (0.060) | **0.246 (0.085)** | **0.331 (0.083)** | **0.474 (0.112)** | **0.473 (0.144)** |
| *RL<sub>accuracy</sub>* | **0.122 (0.060)** | 0.122 (0.061) | 0.055 (0.043) | **0.127 (0.059)** | 0.091 (0.051) | **0.148 (0.062)** | **0.205 (0.056)** |

Among the subset of 6 cognitive performance measures, all estimated among-individual correlations (r_I_) were positive (Table 3). This included among traits in the association task (r*_ALtime. ALspeed_* = 0.758 (0.074), r*_ALtime. ALaccuracy_* = 0.729 (0.092), r*_ALspeed. ALaccuracy_* = 0.894 (0.039)) meaning that we find no support for a speed-accuracy trade-off here. Similarly, both variables in the reversal learning task (r*_RLtime. RLaccuracy_* = 0.473 (0.144)) were positively correlated. There was thus no evidence of negative correlations as predicted by specific trade-offs, either within or between cognitive tasks, and rather the correlation structure of **ID** is uniformly positive, consistent with the *g*-model of general cognitive performance (Table 3).

Eigen decomposition of **ID_rc_** reflects this, with the first major axis (first principle component, PC1, with 95% confidence intervals from 5000 bootstrap replicates) explaining 57.1% (42.7%, 69.8%) of the among-individual variation in multivariate phenotype on a correlation scale. Subsequent vectors necessarily explain sequentially diminishing amounts of variation (PC2 = 25.4% [12.6, 30.9]; PC3 = 12.1% [7.4, 15.6]; PC4 = 3.7% [4.7, 9.9]; PC5 = 1.6% [0.8, 5.5]; PC6 > 0.001% [0,0]). All 6 cognitive performance measures load with the same sign on PC1 and these loadings are statistically significant based on bootstrapped confidence intervals not overlapping zero (Figure 3; Supplementary Information Table S2). Thus, the eigen decomposition reiterates the view that the correlation structure is consistent with a dominant axis of among-individual variation in cognitive phenotype caused by differences in underlying *g-*model of general cognitive performance. On this axis, individuals at one end of the axis can be considered to have ‘higher/better’ performance in all tasks (i.e. are faster and more accurate in the association task, faster to solve the motor learning task, and faster and more accurate in the reversal learning task when the cue is reversed), while individuals at the other end perform relatively poorly in all respects.

**Figure 3:**
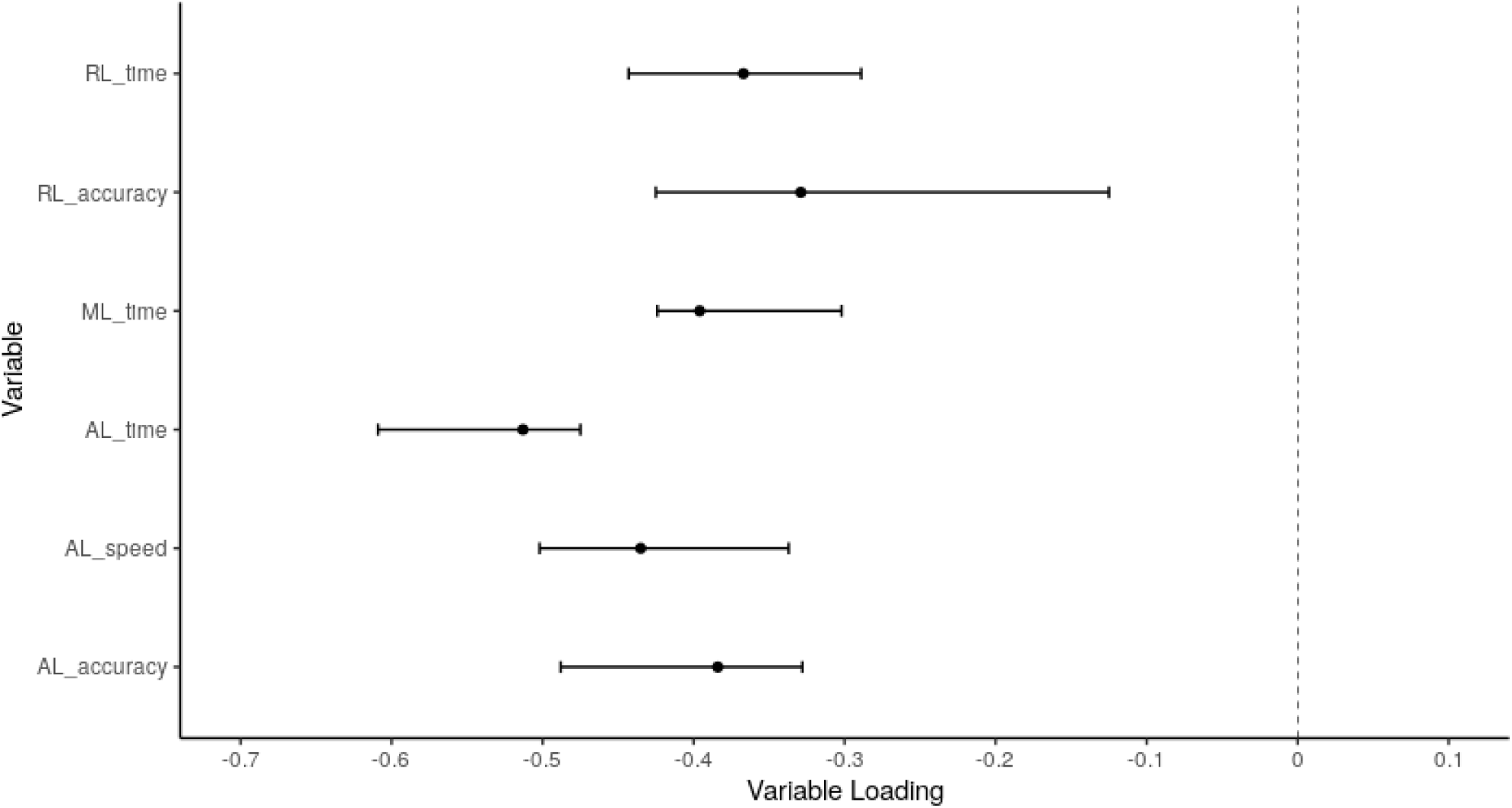
Variable loadings from the first eigen vector (principal component, PC1) of the **ID**-matrix. This axis explains 57.1% of the among-individual (co)variation found in the components of the cognitive phenotype in our guppy population. Points show variable loadings from the first eigen vector of our estimate of **ID**, with bars representing 95% confidence intervals on each loading (calculated from 5000 bootstrapped replicates of the model)

## Discussion

In this study we sought to determine whether – and to what extent – there exists among-individual variation for cognitive performance in a captive population of recently wild-derived guppies. Adopting a multivariate approach allowed us to test for covariation among multiple cognitive performance measures, as well as between cognitive performance and a shy-bold type axis of personality variation. Our study yields three main findings. First, there is among-individual variation underpinning all cognitive performance measures. Second, there is strong among-individual correlation structure among cognitive performance measures and bold type personality. Thirdly, the structure of **ID**-matrix provides no support for the presence of important performance trade-offs, either between cognitive tasks, or between speed and accuracy within learning tasks. Rather we suggest that the matrix is consistent with a single axis of general variation that causes positively correlated performance across all cognitive tasks. Individuals at one end of the axis can be considered to have relatively low ‘general cognitive performance’ compared to those at the other end of the axis. The structure of **ID**-matrix is therefore consistent with expectations under the *g-*model, although it is important to acknowledge that other latent factors could exist that create similar patterns of covariation among observed behaviours. In what follows, we discuss these findings in the context of understanding variation in, and evolution of, animal cognition, while also drawing attention to important assumptions and caveats underpinning our conclusions.

Among-individual variation was found in each of the cognitive performance measures as well as in the measure of personality tested. That guppies show consistent among-individual differences in ‘boldness’, as measured here by time to emerge into the test compartment over the first set of 9 trials (with a conditional repeatability of 43.9%) was expected from earlier studies of this population (e.g. Houslay et al., 2019) and guppies more widely (Brown & Irving, 2014; O’Neill et al., 2018; Gasparini et al., 2019). Indeed similar findings are common across a wide range of fish models used in behavioural studies (Boulton et al., 2018; White et al., 2020). We also find among-individual variation in all measures of cognitive performance across each task. Thus, we conclude that individuals differ in; ability to discriminate between rewarded and unrewarded colours in an association learning task, ability to learn a novel motor learning task, and, in cognitive flexibility as tested by a reversal learning task. The estimated repeatabilities of our observed cognitive performance measures are moderate relative to behavioural traits generally (Bell et al., 2009), but generally consistent with reports for cognitive assays in animals (Cauchoix et al., 2018). It is worth noting that behavioural repeatabilities generally decline as inter-observation period increases (Boulton et al., 2014) and here the inter-observation period was short (usually less than 24hrs with up to 9 observations made in a 48hr period for some traits). Although this is common for cognitive studies, we acknowledge that the stability of individual differences in cognitive performance may well be less over longer timeframes.

Our multivariate modelling provides no support for trade-offs among cognitive performance measures. This conclusion stems from the finding that the correlation structure in **ID** is universally positive. Thus, for example, speed-accuracy trade-offs are not detected in our association learning trials. Speed-accuracy trade-offs are widely predicted to arise where behavioural outcomes depend on cognitive decision making (Briffa, 2013), although this prediction assumes that faster decisions are better (if they come at no cost to accuracy). They have been detected in some animal studies, for instance in Carib grackles (*Quiscalus lugubris*) where individuals that solve novel problems faster also make more errors in discrimination learning tasks (Ducatez et al., 2019). Here we actually find the opposite pattern, faster individuals are also more accurate, a result that seems counterintuitive but is consistent with reports in song sparrows (*Melospiza melodia*; Boogert, 2011), and bumblebees (*Bombus terrestris;* Raine, 2012). We note that the correlation structure of behavioural **ID**-matrices can be sensitive to environmental context (Houslay et al., 2018) as a consequence of IxE interactions (among-individual differences in plasticity; Nussey et al., 2007). It is therefore possible trade-offs could be apparent only under specific conditions and/or assays. This scenario is suggested by a recent study of archerfish in which a speed-accuracy trade-off was only statistically significant under the more challenging of two cognitive testing treatments applied (Jones et al., 2020). Plastic responses to social context could also be important since, for instance, in less competitive scenarios (including the single housing used here) slow decisions may carry minimal costs removing motivation to be fast (Sih & Del Giudice, 2012).

More broadly, we also find positive correlations across cognitive performance variables measured within different cognitive tasks. This is counter to predictions under hypothesised trade-off models for maintaining variation. For instance, it has been hypothesised that individuals more reactive to changes in environmental cues will perform relatively poorly in association learning but –being cognitively flexible – will do well in reversal learning tasks (Sih & Del Giudice, 2012; Bebus et al., 2016). Here we find a strong positive among-individual correlation between our measures of performance in association and reversal learning (r_I_=0.54). Again we note the absence of apparent trade-offs has precedent in the animal cognition literature. For example, positive correlations have been reported between discrimination acquisition and reversal learning in sparrows (Boogert et al., 2011), social learning and innovative problem solving in pigeons (*Columba livia*; Bouchard et al., 2007), and motor skill learning and reversal learning in pheasants (van Horik et al., 2018).

If positive correlations in the **ID**-matrix provide no support for trade-offs, then the corollary is that they are consistent with a general axis of cognitive variation among-individuals in this population. Thus, excluding the boldness proxy of emergence time and focusing on cognitive performance measures only, we found that 57% of the variation in correlation matrix was explained by the leading principle component (or eigen vector). By comparison, in tests of the general intelligence model of cognition using human psychometric test batteries, an underlying *g-*factor is typically found to account for about 40–50% of the multivariate variation (Carroll, 1993; Deary, 2001). Under the *g-*model, trait performance in cognitive assays across multiple domains are expected to load strongly (and in the same direction) on a dominant first principle component (Plomin, 2001; Plomin & Spinath, 2002). We note that human psychometric studies usually test a different and far wider range of cognitive abilities than examined here. Nonetheless, here we find a similar pattern of strong positive loadings on a dominant first principle component. Our results thus add to a small, but growing body of empirical literature finding support for general axis of variation among-individuals in animal cognitive ability (for example see Ashton et al., 2018; Shaw et al., 2015).

Our conclusion that (multivariate) cognitive variation in guppies is consistent with an underlying general cognitive performance factor requires some caveats. These largely arise because latent variables are, by definition, unmeasurable. Thus, all inferences about (latent) cognitive traits from (observed) behavioural performance measures are inevitably assumption-laden. First, we are limited by the set of variables assayed. Inclusion of multiple cognitive domains, different assays, and/or different contexts, would provide more robust insight into the biological extent and evolutionary potential of general intelligence (Poirier et al., 2020). Thus, for instance, we detect no evidence of cognitive trade-offs within performance variable measured within each cognitive task, but they could occur among a wider set of traits (or contexts) not examined. Here we used three cognitive tasks as proxies of three cognitive domains: association learning; motor learning; and reversal learning, but did so using a single testing paradigm in which performance measures share a requirement for individuals to discriminate between two coloured cues. In this way, cognitive performance measures across all three tasks could be said to be based on common associative processes, resulting ultimately in a predisposition for finding a positive correlation structure among them. Second, the eigen vector decomposition provides only a statistical description of the estimated correlation structure among observed variables; that **ID** is consistent with an axis of variation in an underlying general cognitive performance factor does not prove such a factor exists. Third, the positive correlation structure could be explained, at least in part, by shared dependence of the observed traits on some other parameter that varies among individuals such as metabolic state (Biro & Stamps, 2010; Mathot & Dall, 2013; McKenzie et al., 2016), satiety (Shettleworth, 1972; Ben-Shahar & Robinson, 2001), or other drivers of motivation (van Horik & Madden, 2016). Among-individual differences could result from differences in motivation rather than differences in cognitive ability per se. For example, the increase in mean performance time in the association task could be confounded by changes in motivation due to trial fatigue (reduced motivation), at least for some individuals, even if some individuals clearly di<u>d</u> perform better over time. Experimental conditions can have important consequences for robust interpretation of studies targeting cognitive traits (Rowe & Healy, 2014; Griffin et al., 2015; Boogert et al., 2018) and while we attempted to minimise ‘non-target’ sources of among-individual variation (e.g. by standardising food rations, housing and water chemistry) we cannot exclude the possibility that these contribute to the structure of **ID**. Finally, we acknowledge the potential for bias in our findings caused by selective removal of individuals due to our imposed learning criterion. Failure to participate in a cognitive task may result in sampling biases when measuring among-individual variation in cognitive performances (Thornton & Lukas, 2012; van Horik et al., 2017). Our learning criterion was specifically weak so that all included individuals represented, as far as possible, among-individual variation of the initial population.

We also found that all six cognitive performance measures were positively correlated with ‘boldness’, measured here as an individual’s tendency to consistently emerge rapidly from the home compartment during trial observations. Pairwise correlations were nominally significant in four of the six cases. Our finding that bolder individuals perform better in cognitive tasks mirrors results reported from several other studies. For instance, boldness has been positively linked with performance speed in associative learning tasks in rainbow trout (*Oncorhynchus mykiss*; Sneddon, 2003), shape discrimination in Cavies (*Cavia aperea*; Guenther et al., 2014), and reversal learning in the Chimango Caracara, (*Milvago chimango;* Guido et al., 2017). While shy-bold type behavioural variation can have important fitness consequences (Wilson et al., 2010; Ariyomo & Watt, 2012; Ballew et al., 2017), estimates of contemporary selection on boldness in wild guppies are lacking. Nonetheless, multiple studies have reported among-population differences in boldness mapping to differences in predation, a link that is consistent with the hypothesis that variable levels of risk could impose balancing selection that maintains variation in boldness. Very speculatively, if this is true, and if correlations in **ID** are recapitulated in **G,** the link between boldness and cognitive performance could help maintain variation in the latter. This is because selection on any one trait (e.g. boldness) has consequences for the evolutionary dynamics of genetically correlated traits (Roff, 2002). We do know that shy-bold type personality variation is heritable in the population of guppies under lab conditions, (White & Wilson, 2019; White et al., 2019; Prentice et al., 2020) but not how selection is acting in the field.

In summary, we find evidence that individual guppies do differ in cognitive performance as measured by a testing protocol that targets association learning, motor learning and cognitive flexibility (reversal learning). We find no support for trade-offs occurring either between speed and accuracy (e.g. within the association learning and reversal learning trials) or between overall performance across cognitive tasks. Subject to the caveats highlighted above, the absence of trade-offs and the presence of strong positive general correlations in **ID** mean that the general intelligence model provides a good description of the structure of multivariate general cognitive performance variation in this species. Our results also suggest that, to the extent that directional natural selection favours higher performance in any single cognitive performance measure, it would lead to positively correlated evolutionary responses across all other measures. However, this inference is dependent on **ID** being a reliable proxy for the **G** matrix (Brommer, 2013). Greater efforts to quantify **G** is thus an obvious, if challenging, next step towards understanding the maintenance of variation in and evolutionary dynamics of animal cognition.

