## Supplemental Information for "A multivariate view of cognitive performance reveals positive correlation in the Trinidadian Guppy (*Poecilia reticulata*)"

**Table S1: Fixed effects estimates for the full univariate models for each variable**

**Table S2: Principle components loadings (eigen decomposition) of ID matrix**

**Figure S1: Plots of raw data of all variables across each set of trials**

**Table S1 – Fixed effects estimates for the full univariate models** Fixed effect estimates (with standard errors in parentheses) from full univariate models for each variable. Intercept represents estimate at Trial number = 1, sex = Female, Stack = A, Colour = Blue and Reward side = Left.

**Table S1a** – *emergence*

| Variable | | Fixed effect | Effect size (SE) | DF | F | P |
|---|---|---|---|---|---|---|
| *emergence* | Mean | Intercept | 0.555 (0.213) | 1, 81.9 | 0.002 | 0.966 |
| | | Trial number (2) | -0.082 (0.136) | 8, 555.7 | 9.240 | <0.001 |
| | | Trial number (3) | 0.425 (0.138) | | | |
| | | Trial number (4) | 0.580 (0.138) | | | |
| | | Trial number (5) | 0.475 (0.118) | | | |
| | | Trial number (6) | 0.523 (0.125) | | | |
| | | Trial number (7) | 0.572 (0.121) | | | |
| | | Trial number (8) | 0.603 (0.126) | | | |
| | | Trial number (9) | 0.363 (0.125) | | | |

| | | | | | |
|---|---|---|---|---|---|
| | Sex (Male) | -0.324 (0.168) | 1, 82.4 | 3.705 | 0.057 |
| | Stack (B) | 0.101 (0.168) | 1, 82.6 | 2.328 | 0.550 |
| | Colour (Green) | -0.030 (0.152) | 1, 81.9 | 0.034 | 0.846 |
| | Reward side (Right) | -0.113 (0.070) | 1, 561.8 | 2.585 | 0.108 |

**Table S1b** – $AL_{speed}$

| Variable | | Fixed effect | Effect size (SE) | DF | F | P |
|---|---|---|---|---|---|---|
| $AL_{speed}$ | Mean | Intercept | -0.657 (0.216) | 1, 81.7 | 0.002 | 0.970 |
| | | Trial number (2) | 0.131 (0.134) | 8, 555.0 | 7.380 | <0.001 |
| | | Trial number (3) | 0.423 (0.136) | | | |
| | | Trial number (4) | 0.659 (0.137) | | | |
| | | Trial number (5) | 0.499 (0.117) | | | |
| | | Trial number (6) | 0.648 (0.124) | | | |
| | | Trial number (7) | 0.631 (0.120) | | | |
| | | Trial number (8) | 0.508 (0.124) | | | |
| | | Trial number (9) | 0.395 (0.216) | | | |
| | | Sex (Male) | 0.197 (0.172) | 1, 82.3 | 1.347 | 0.261 |
| | | Stack (B) | -0.102 (0.172) | 1, 82.4 | 1.277 | 0.554 |
| | | Colour (Green) | 0.223 (0.155) | 1, 81.7 | 2.011 | 0.156 |
| | | Reward side (Right) | 0.148 (0.070) | 1, 560.9 | 4.552 | 0.033 |

**Table S1c** – $AL_{accuracy}$

| Variable | | Fixed effect | Effect size (SE) | DF | F | P |
|---|---|---|---|---|---|---|
| $AL_{accuracy}$ | Mean | Intercept | -0.645 (0.177) | 1, 79.3 | 0.002 | 0.963 |
| | | Trial number (2) | -0.137 (0.140) | 8, 558.0 | 4.602 | <0.001 |
| | | Trial number (3) | 0.203 (0.142) | | | |
| | | Trial number (4) | 0.330 (0.142) | | | |
| | | Trial number (5) | 0.361 (0.122) | | | |
| | | Trial number (6) | 0.374 (0.124) | | | |
| | | Trial number (7) | 0.395 (0.124) | | | |
| | | Trial number (8) | 0.308 (0.129) | | | |
| | | Trial number (9) | 0.230 (0.177) | | | |
| | | Sex (Male) | 0.103 (0.127) | 1, 79.3 | 1.049 | 0.418 |
| | | Stack (B) | -0.205 (0.127) | 1, 79.5 | 4.229 | 0.112 |
| | | Colour (Green) | 0.907 (0.115) | 1. 79.3 | 61.770 | <0.001 |
| | | Reward side (Right) | 0.030 (0.072) | 1, 567.4 | 0.161 | 0.689 |

**Table S1d** – $AL_{time}$

| Variable | | Fixed effect | Effect size (SE) | DF | F | P |
|---|---|---|---|---|---|---|
| $AL_{time}$ | Mean | Intercept | 0.151 (0.227) | 1, 58.5 | 0.146 | 0.704 |
| | | Trial number (2) | -0.224 (0.149) | 17, 801.8 | 2.246 | 0.002 |
| | | Trial number (3) | -0.250 (0.147) | | | |
| | | Trial number (4) | -0.043 (0.154) | | | |
| | | Trial number (5) | -0.298 (0.150) | | | |
| | | Trial number (6) | -0.313 (0.174) | | | |
| | | Trial number (7) | -0.265 (0.170) | | | |
| | | Trial number (8) | -0.417 (0.158) | | | |
| | | Trial number (9) | -0.403 (0.149) | | | |
| | | Trial number (10) | -0.377 (0.152) | | | |
| | | Trial number (11) | -0.567 (0.154) | | | |
| | | Trial number (12) | -0.269 (0.176) | | | |
| | | Trial number (13) | -0.414 (0.158) | | | |
| | | Trial number (14) | -0.527 (0.155) | | | |
| | | Trial number (15) | -0.587 (0.178) | | | |
| | | Trial number (16) | -0.512 (0.154) | | | |
| | | Trial number (17) | -0.688 (0.182) | | | |
| | | Trial number (18) | -0.553 (0.227) | | | |

| | | | | | |
|---|---|---|---|---|---|
| | Sex (Male) | 0.315 (0.181) | 1, 59.5 | 2.381 | 0.087 |
| | Stack (B) | 0.430 (0.181) | 1, 59.2 | 3.606 | 0.021 |
| | Colour (Green) | -0.362 (0.146) | 1, 87.8 | 5.894 | 0.015 |
| | Reward side (Right) | -0.095 (0.065) | 1, 803.4 | 2.110 | 0.147 |

**Table S1e** – $ML_{time}$

| Variable | | Fixed effect | Effect size (SE) | DF | F | P |
|---|---|---|---|---|---|---|
| $ML_{time}$ | Mean | Intercept | 0.242 (0.244) | 1, 48.1 | 0.002 | 0.967 |
| | | Trial number (2) | 0.028 (0.174) | 17, 692.6 | 1.174 | <0.001 |
| | | Trial number (3) | -0.099 (0.174) | | | |
| | | Trial number (4) | 0.143 (0.149) | | | |
| | | Trial number (5) | -0.288 (0.171) | | | |
| | | Trial number (6) | 0.178 (0.154) | | | |
| | | Trial number (7) | -0.037 (0.171) | | | |
| | | Trial number (8) | 0.386 (0.174) | | | |
| | | Trial number (9) | 0.104 (0.172) | | | |
| | | Trial number (10) | -0.332 (0.147) | | | |
| | | Trial number (11) | -0.516 (0.229) | | | |
| | | Trial number (12) | -0.468 (0.150) | | | |
| | | Trial number (13) | -0.309 (0.151) | | | |
| | | Trial number (14) | -0.272 (0.152) | | | |
| | | Trial number (15) | -0.452 (0.148) | | | |
| | | Trial number (16) | -0.166 (0.149) | | | |
| | | Trial number (17) | -0.233 (0.152) | | | |
| | | Trial number (18) | -0.502 (0.146) | | | |

| | | | | | |
|---|---|---|---|---|---|
| | Sex (Male) | -0.063 (0.192) | 1, 48.3 | -0.974 | 0.742 |
| | Stack (B) | 0.262 (0.193) | 1, 48.3 | 3.742 | 0.180 |
| | Colour (Green) | -0.560 (0.158) | 1, 48.1 | 12.61 | 0.001 |
| | Reward side (Right) | 0.279 (0.072) | 1, 694.8 | 15.07 | <0.001 |

**Table S1f** – RL$_{time}$

| Variable | | Fixed effect | Effect size (SE) | DF | F | P |
|---|---|---|---|---|---|---|
| $RL_{time}$ | Mean | Intercept | -0.341 (0.307) | 1, 47.7 | 0.005 | 0.946 |
| | | Trial number (2) | 0.196 (0.197) | 17, 581.2 | 1.565 | 0.060 |
| | | Trial number (3) | 0.272 (0.214) | | | |
| | | Trial number (4) | 0.328 (0.195) | | | |
| | | Trial number (5) | -0.012 (0.193) | | | |
| | | Trial number (6) | 0.332 (0.190) | | | |
| | | Trial number (7) | 0.224 (0.197) | | | |
| | | Trial number (8) | 0.360 (0.196) | | | |
| | | Trial number (9) | 0.344 (0.213) | | | |
| | | Trial number (10) | 1.132 (0.594) | | | |
| | | Trial number (11) | 0.343 (0.234) | | | |
| | | Trial number (12) | 0.449 (0.218) | | | |
| | | Trial number (13) | 0.272 (0.205) | | | |
| | | Trial number (14) | 0.487 (0.254) | | | |
| | | Trial number (15) | 0.494 (0.206) | | | |
| | | Trial number (16) | 0.616 (0.200) | | | |
| | | Trial number (17) | 0.503 (0.206) | | | |
| | | Trial number (18) | 0.310 (0.205) | | | |

| | | | | | |
|---|---|---|---|---|---|
| | Sex (Male) | -0.104 (0.219) | 1, 47.7 | 0.169 | 0.638 |
| | Stack (B) | -0.153 (0.227) | 1, 47.9 | 0.380 | 0.505 |
| | Colour (Green) | 0.348 (0.182) | 1, 47.4 | 3.529 | 0.066 |
| | Reward side (Right) | 0.043 (0.078) | 1, 578.4 | 0.308 | 0.579 |

**Table S1g** – $RL_{accuracy}$

| Variable | | Fixed effect | Effect size (SE) | DF | F | P |
|---|---|---|---|---|---|---|
| $RL_{accuracy}$ | Mean | Intercept | -0.402 (0.263) | 1, 46.3 | 0.110 | 0.742 |
| | | Trial number (2) | 0.206 (0.211) | 17, 584.2 | 1.085 | 0.271 |
| | | Trial number (3) | 0.104 (0.230) | | | |
| | | Trial number (4) | 0.350 (0.209) | | | |
| | | Trial number (5) | 0.280 (0.206) | | | |
| | | Trial number (6) | 0.290 (0.203) | | | |
| | | Trial number (7) | 0.279 (0.211) | | | |
| | | Trial number (8) | 0.185 (0.210) | | | |
| | | Trial number (9) | 0.315 (0.228) | | | |
| | | Trial number (10) | 0.058 (0.636) | | | |
| | | Trial number (11) | 0.001 (0.260) | | | |
| | | Trial number (12) | 0.327 (0.233) | | | |
| | | Trial number (13) | 0.359 (0.219) | | | |
| | | Trial number (14) | 0.225 (0.272) | | | |
| | | Trial number (15) | 0.548 (0.220) | | | |
| | | Trial number (16) | 0.514 (0.214) | | | |
| | | Trial number (17) | 0.610 (0.220) | | | |
| | | Trial number (18) | 0.437 (0.219) | | | |

| | | | | |
|---|---|---|---|---|
| Sex (Male) | -0.008 (0.166) | 1, 46.4 | 0.006 | 0.963 |
| Stack (B) | -0.303 (0.173) | 1, 48.1 | 5.464 | 0.086 |
| Colour (Green) | 0.631 (0.138) | 1, 46.1 | 20.860 | <0.001 |
| Reward side (Right) | 0.025 (0.084) | 1, 579.9 | 0.089 | 0.764 |

**S2 – Principle components analyses (eigen decomposition) of ID matrix**

**Table S2a**: Phenotypic variance (%) explained by six principle components of ID (on a correlation scale), with 95% confidence intervals from 5000 bootstrap replicates

| PC1 | PC2 | PC3 | PC4 | PC5 | PC6 |
|---|---|---|---|---|---|
| 0.571 [0.427, 0.698] | 0.254 [0.126, 0.309] | 0.121 [0.074, 0.156] | 0.037 [0.047, 0.099] | 0.017 [0.008, 0.055] | 0.001 [0, 0] |

**Table S2b:** Loadings of six cognitive variables onto six principle components of ID (on a correlation scale)

| | PC1 | PC2 | PC3 | PC4 | PC5 | PC6 |
|---|---|---|---|---|---|---|
| $AL_{speed}$ | -0.435 | 0.431 | -0.022 | -0.217 | 0.758 | -0.047 |
| $AL_{accuracy}$ | -0.384 | 0.543 | 0.081 | -0.198 | -0.555 | 0.451 |
| $AL_{time}$ | -0.513 | 0.056 | 0.254 | 0.423 | -0.240 | -0.658 |
| $ML_{time}$ | -0.396 | -0.471 | 0.343 | 0.373 | 0.193 | 0.571 |
| $RL_{time}$ | -0.367 | -0.523 | 0.022 | -0.737 | -0.135 | -0.174 |
| $RL_{accuracy}$ | -0.329 | -0.139 | -0.900 | 0.231 | -0.065 | 0.070 |

**Figure S1a**:

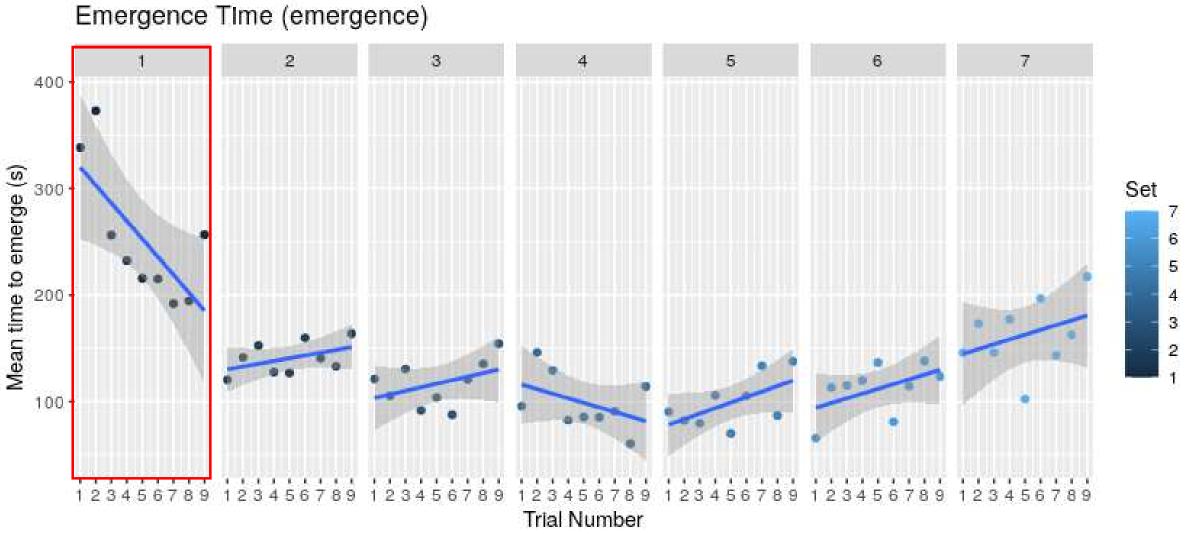


**Figure S1b**:

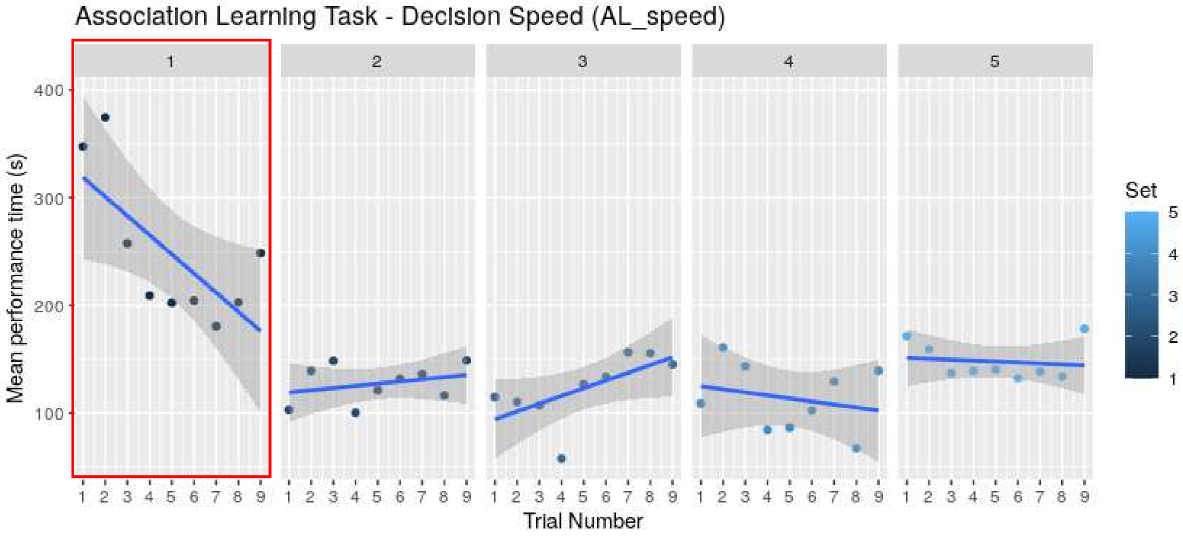

**Figure S1c**:

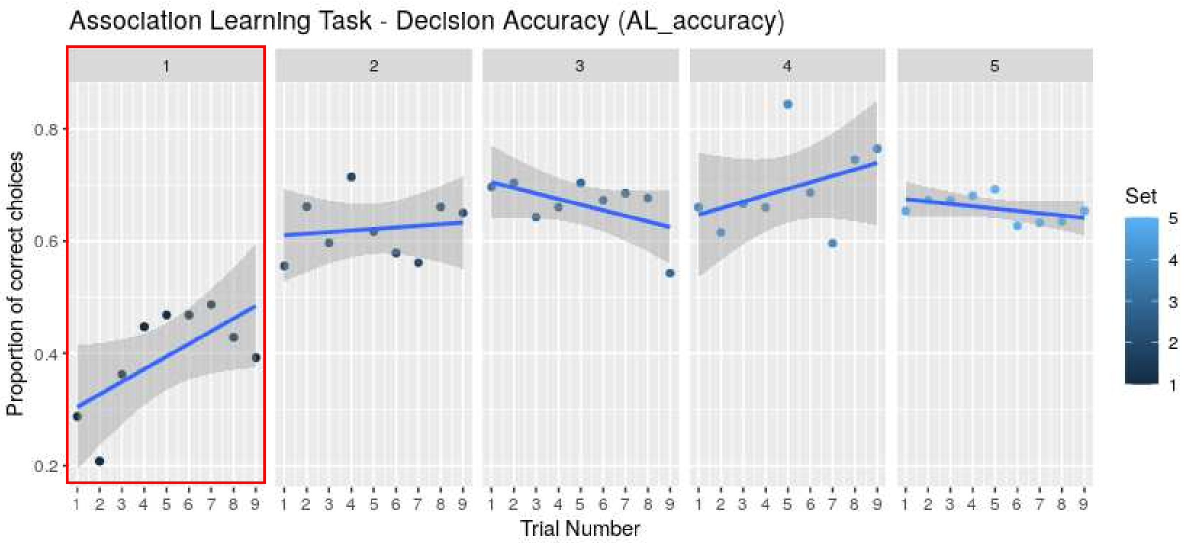


**Figure S1d**:

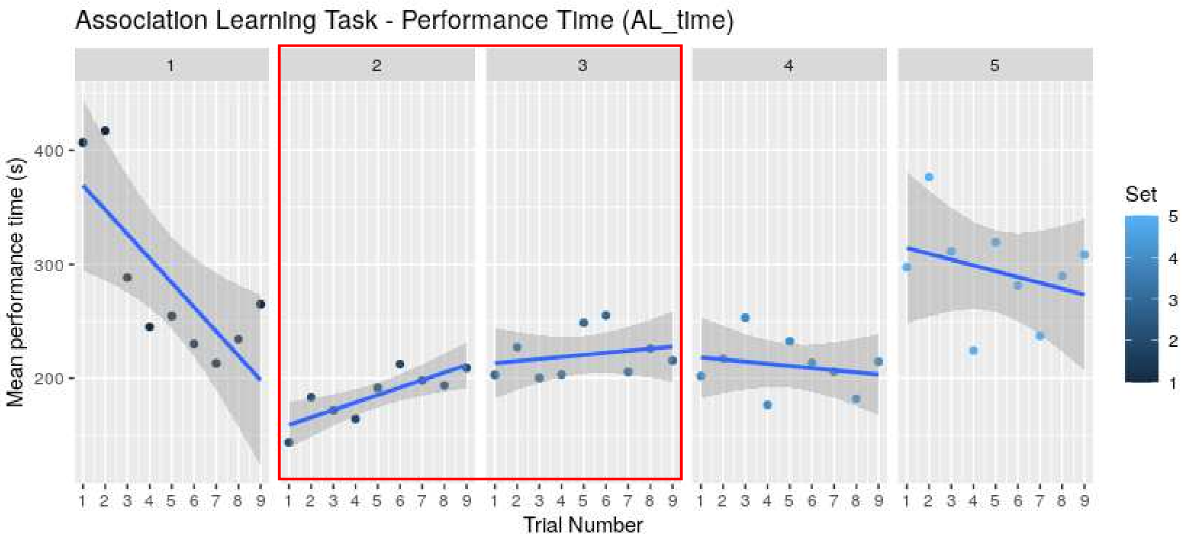

**Figure S1e**:

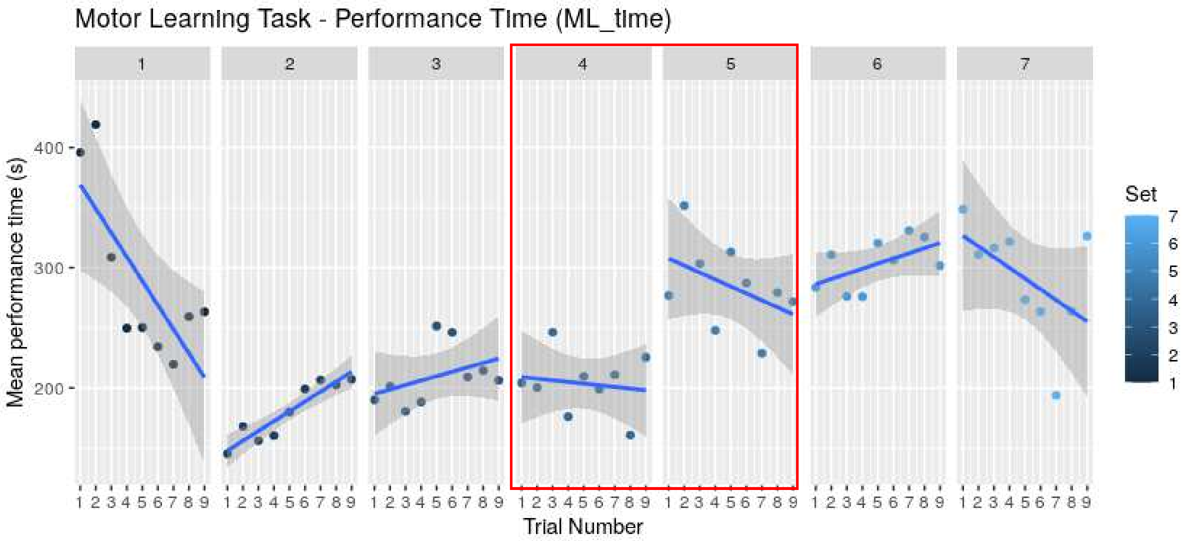


**Figure S1f**:

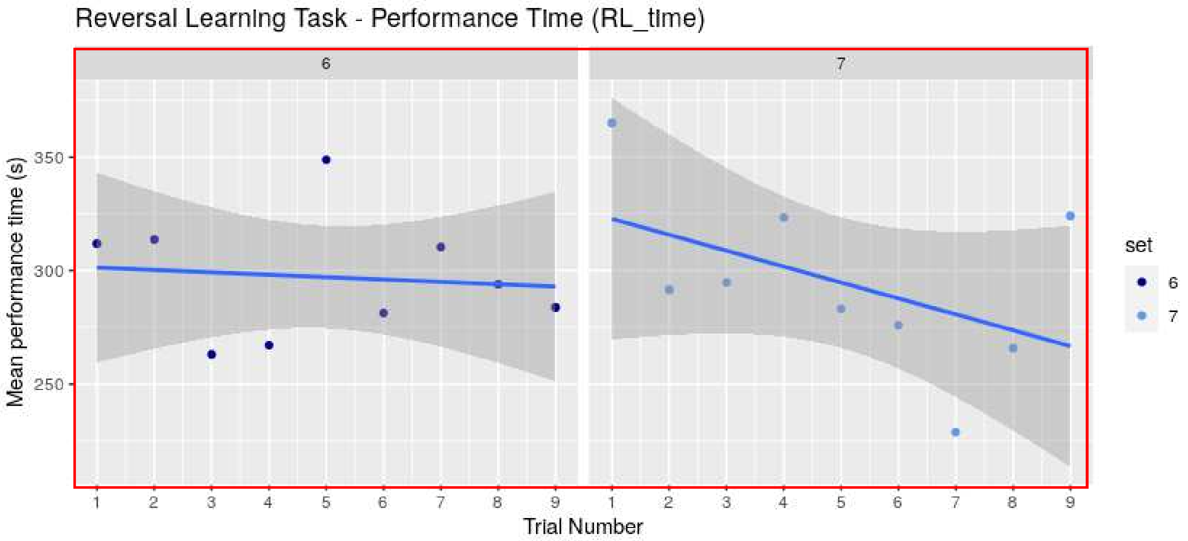

**Figure S1g**:

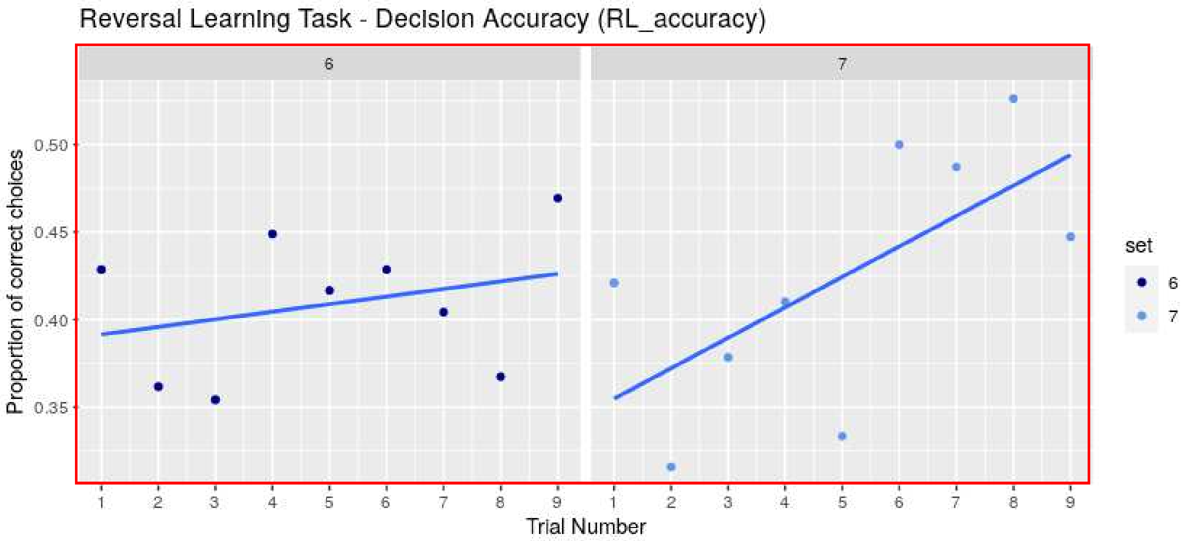


**Figure S1.** Plots of observed means for the personality trait and all the cognitive performance measures across each set of 9 trials. Data are shown from all trials to visualise overall patterns, but red boxes denote informative trials in which observations were used as proxies for a) emergence time (*emergence)*, the association learning task b) $AL_{speed}$, c) $AL_{accuracy}$, d) $AL_{time}$, the motor learning task e) $ML_{time}$, and reversal learning task f) $RL_{time}$ and g) $RL_{accuracy}$. Blue points denote means at each trial number within each set, and line show regressions through these means.
